# Boldine prevents diabetes-induced skeletal muscle dysfunction by inhibiting large-pore channels

**DOI:** 10.64898/2026.02.24.707704

**Authors:** Walter Vásquez, Luis A. Cea, Felipe Troncoso, Hermes Sandoval, Andrea Lira, Xavier F. Figueroa, Carlos Escudero, Juan C. Sáez

**Affiliations:** Facultad de Ciencias Biológicas, Pontificia Universidad Católica de Chile, Santiago; Instituto de Ciencias Biomédicas, Facultad de Ciencias de la Salud, Universidad Autónoma de Chile, Santiago, Chile; Laboratorio de Fisiología Vascular, Group of Research and Innovation in Vascular Health, GRIVAS Health, Departamento de Ciencias, Universidad del Bio-Bio, Chile; Departamento de Morfología, Facultad de Medicina, Universidad Andres Bello, Santiago, Chile; Consortium of Neurovascular Research and Innovation (NEUROVAS), Chillán, Chile; Instituto de Neurociencias, Centro Interdisciplinario de Neurociencias de Valparaíso, Universidad de Valparaíso, Valparaíso, Chile

**Keywords:** PPARγ, lipid accumulation, inflammation, myopathy, sarcolemma permeability, hemichannel blocker

## Abstract

**Background:** Diabetes mellitus leads to skeletal muscle dysfunction associated with loss of strength, impaired blood perfusion, lipid accumulation, and inflammation. The opening of large-pore channels has been linked to increased membrane permeability and inflammatory signaling in several pathologies. Boldine, an alkaloid from *Peumus boldus*, blocks large-pore channel activity and exhibits antioxidant and anti-inflammatory properties. This study evaluated whether boldine prevents skeletal muscle alterations induced by diabetes and explored potential underlying mechanisms.

**Methods:** Diabetes was induced in male C57BL/6J mice using streptozotocin (STZ, 40 mg/kg/day for 5 days). Diabetic mice were treated with boldine (50 mg/kg/day) for four weeks. Muscle strength and resting membrane potential were analyzed in vivo. Also, right gastrocnemius muscle blood perfusion at basal and after acetylcholine (10 μM) stimulation were analyzed in vivo. Lipid accumulation was assessed using Oil Red O staining, and CD31 immunodetection was used to evaluate capillary density. mRNA levels of *NLRP3* were evaluated in muscle by qPCR. In human myoblasts (AB1167) cultured under low (8 mM) or high glucose (25 mM) conditions, with or without boldine, membrane permeability (ethidium uptake), intracellular Ca²⁺ (Fura-2), nitric oxide (DAF-FM), and levels of *NLRP3* and *Casp1* (qPCR) and reactivity *PPARγ* (Immunofluorescence) were determined.

**Results:** STZ mice showed reduced muscle strength and depolarized resting membrane potential, both prevented by boldine. Basal muscle perfusion was ∼20% lower in diabetic mice (160.1 ± 17.2 vs. 199.1 ± 13.8 units), whereas boldine preserved perfusion (184.6 ± 14.3 units). Oil Red O–positive fibers increased to 52.4 ± 3.6% in diabetic mice and decreased to 15.2 ± 4.1% with boldine (control: 3.1 ± 1.3%; p<0.05). NLRP3 mRNA increased 17.7 ± 2.8-fold in diabetic muscle and was reduced by ∼50% with boldine. In myoblasts, high glucose increased ethidium uptake, nitric oxide production, NLRP3 and caspase-1 expression, and nuclear PPARγ (∼45% positive nuclei); all effects were prevented by boldine.

**Conclusions:** Boldine preserves skeletal muscle function and vascular reactivity in diabetes and prevents lipid accumulation and inflammasome activation both in vivo and in vitro. These effects are associated with inhibition of large-pore channel activity and attenuation of downstream calcium-dependent, inflammatory, and adipogenic pathways, supporting boldine as a promising therapeutic candidate for diabetes-associated skeletal muscle dysfunction.

**Graphical abstract:** In myoblasts, high glucose activates large-pore channels, elevating cytoplasmic Ca²⁺ concentration and nitric oxide generation, which increases the activity of Cx-formed hemichannels, raises the levels of inflammasome components, and promotes lipid accumulation. In STZ-diabetic mice, de novo expression of large-pore channels in skeletal muscles contributes to reduced blood perfusion, accumulation of intramuscular fat, muscle weakness, and reduced resting membrane potential of myofibers. Boldine inhibits large-pore channel activity, preventing these alterations and preserving muscle physiology *in vivo*.

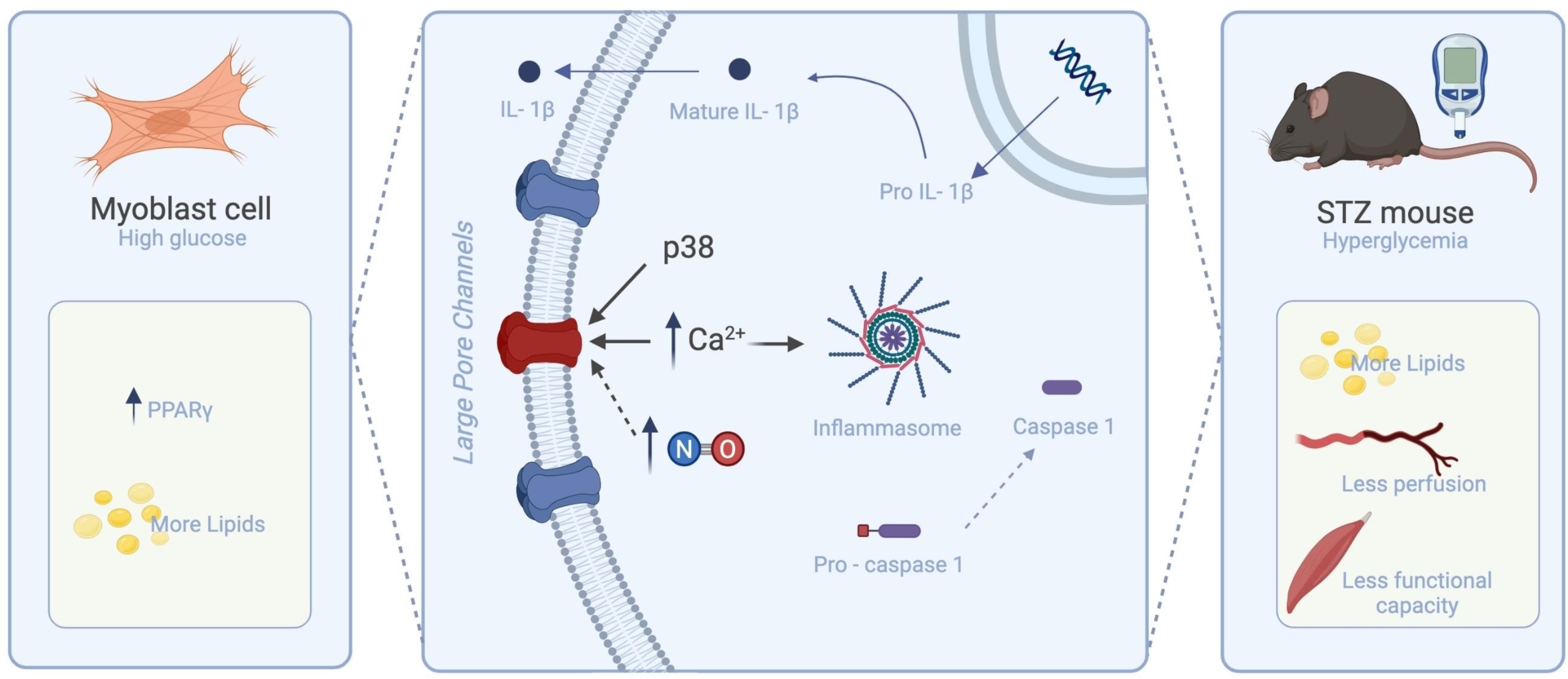

## INTRODUCTION

Diabetes mellitus (DM) is a globally prevalent chronic disease, characterized by insufficient insulin production or reduced capacity for its effective use [1]. One significant complication of DM is skeletal muscle myopathy (SMM). SMM is particularly relevant as skeletal muscle is crucial for glucose uptake and its deterioration profoundly impacts whole-body glucose homeostasis [2]. SMM manifests as muscle atrophy, decreased regeneration capacity, and fat infiltration, with adult skeletal muscle satellite cells (muscle progenitors) being especially vulnerable to the diabetic environment [3].

Under normal conditions, satellite cells remain quiescent, but upon stimuli like injury, they proliferate into myoblasts committed to the myogenic lineage, eventually fusing into myotubes and mature myofibers [4]. However, high glucose exposure promotes adipogenic differentiation in skeletal muscle-derived stem cells, leading to ectopic intramuscular fat accumulation [5]. Recent studies also implicate fibroadipogenic progenitors in this process, though the underlying molecular mechanisms are still poorly understood [6].

Notably, inactivation of connexin 43 (Cx43) and Cx45 expression in skeletal myofibers prevented lipid accumulation in a murine dysferlinopathy model, suggesting a key role for connexin-formed hemichannels in muscle fat infiltration and dysfunction [7]. Connexins (Cxs) and pannexin1 (Panx1) form large-pore channels called hemichannels in the plasma membrane [8]. Cx39, Cx43, and Cx45 are expressed in skeletal muscle during progenitor cell proliferation and muscle fiber formation [9], and their aberrant expression or upregulation of Panx1 hemichannels (Panx1 HCs) contributes to various neuromuscular diseases [10].

Boldine, an alkaloid from *Peumus boldus*, is known to block three large-pore channels [11], and previous work has shown it restores normal differentiation and improves muscle function in a dysferlinopathy model [12]. Similar protection has been observed in mice with dysferlinopathy treated with pulverized boldo leaves [13].

In this study, we propose that large-pore channels are key elements in the pathological mechanism of SMM in streptozotocin-induced diabetes and in high glucose-induced aberrant adipogenic commitment during differentiation. We investigated whether boldine could prevent these diabetic myopathy features, including lipid accumulation, impaired muscle function, and local inflammation, thereby preserving skeletal muscle integrity in diabetic animals. These experiments aim to advance in elucidating the therapeutic potential of boldine in diabetes-associated skeletal muscle dysfunction.

## Materials and methods

*Reagents*. Dulbecco’s modified Eagle medium (DMEM), ethidium bromide (Etd^+^), 4-Bromo A23187 were acquired from Sigma-Aldrich (St. Louis, MO, USA), dithiothreitol (DTT), Fura-2AM, DAF-FM, SB203580, A740003, oil Red O, horse serum, and anti-PPAR-γ were purchased from Thermo Fisher Scientific (Waltham, MA, USA). Cy2- and Cy3-conjugated goat anti-rabbit IgG were purchased from Jackson ImmunoResearch Laboratories (West Grove, PA, USA). The hydrochloride form of boldine was prepared as described previously [11]. Streptozotocin (STZ) and acetylcholine was purchased from Sigma-Aldrich (St. Louis, MO, USA). (R)-2-(4-chlorophenyl)-2-oxo-1-phenylethyl quinoline-2-carboxylate (C_24_H_16_ClNO_3_; MW 401.85 g.mol-1), called D4, was synthetized by Edelris s.a.s. Medicinal Keymistry (Lyon, France).

### Animals

Male wildtype (WT) C57Bl/6 mice were used. Diabetes was induced after a 6-hour fasting period, followed by a single intraperitoneal injection of STZ, 40 mg/kg/day for 5 days. To confirm the development of diabetes, blood samples were collected from the tail two days after STZ administration. Mice were considered diabetic if their blood glucose levels were ≥180 mg/dL. Glycaemia was measured using a portable glucometer (Accutrend sensor, Roche, Basel, Switzerland). All protocols were approved by the Bioethics Committee of the Universidad de Valparaíso (CBC 85-2023) in accordance with the ethical standards established in the 1964 Declaration of Helsinki and its later amendments. All efforts were made to minimize animal suffering and to reduce the number of animals used, and alternatives *in vivo* techniques were implemented when it was possible. Three experimental groups were included: Control (n = 4), STZ-induced diabetes (STZ, n = 4), and STZ-induced diabetes treated with boldine (STZ + Boldine, n = 4). Diabetes was induced by intraperitoneal injections of streptozotocin (40 mg/kg/day for 5 consecutive days). Boldine was administered orally at a dose of 50 mg/kg/day for four weeks by mixing the compound with peanut butter, which was voluntarily consumed by the animals. Control and STZ mice received peanut butter without boldine as vehicle control. All experimental groups were euthanized at the same age to avoid age-related confounding effects.

### Forelimb muscle strength

Muscle strength was assessed using a digital force transducer (GPM-100; Melquest, Toyama, Japan) with a triangular metal bar. Mice grasped the bar with their forepaws and were pulled backward until release. The peak force was recorded. Each mouse underwent three consecutive trials per session, with the highest values used for analysis. Measurements were performed at the end of the treatment period in the three experimental groups (Control, STZ, and STZ + Boldine). All measurements were performed head-to-head by a group-blinded experimenter.

### Blood perfusion

Muscle blood flow was assessed *in vivo* using Laser Speckle Contrast Imaging (LSCI) technology (Pericam® PSI-HR system, Perimed, Stockholm, Sweeden). Adult mice were anesthetized with 4% isoflurane. The skin over the right gastrocnemius muscle was removed, and microvascular blood flow was recorded from four predefined circular regions of interest (ROIs). The protocol included a 3–4 min baseline recording, followed by a bolus of acetylcholine (10 µM) to induce endothelium-dependent vasodilation. Perfusion was monitored continuously, and time-of-interest windows (40 seconds) were extracted for basal and post-acetylcholine conditions.

Data acquisition and analysis were conducted by two independent investigators (FT and HS). To minimize bias, experiments were performed in a randomized, head-to-head fashion, simultaneously including mice from all experimental groups. Data analysis was performed in a blinded manner. Changes in blood flow are reported in raw perfusion units as indicated previously [13].

### Isolation of mouse skeletal myofibers

Myofibers were isolated from flexor digitorum brevis (FDB) muscles as previously described. FDB muscles were dissected, incubated in 0.2% collagenase type I, and then gently triturated to disperse single myofibers. Dissociated myofibers were washed and suspended in Krebs solution (in mM: 145 NaCl, 5 KCl, 3 CaCl_2_, 1 MgCl_2_, 5.6 glucose, 10 HEPES-Na, pH 7.4) with 10 μM N-benzyl-p-toluene sulphonamide (BTS) to inhibit contractions and reduce damage.

### Resting membrane potential (RMP)

RMP was recorded from myofibers of the flexor digitorum brevis muscles in a whole-cell configuration at 25°C. RMP recording was performed on fibers kept in culture for maximum 48 h, before the RMP drops as consequence of denervation [14]. The pipette was a borosilicate electrode filled with 3 M KCl, and the bath solution was a Krebs-bicarbonate buffered solution at pH 7.4. The pipette resistance was approximately 50 Ω. All experiments were performed using an Olympus IX 51 inverted microscope, the Axopatch 1-D amplifier, and the Digidata 1322 digitizer and Clampex 9.1 acquisition programs were used. Data were analyzed using Clampfit 2.1.

### Myoblast cultures

The human myoblast cell line (AB1167) previously described [12] were cultured in DMEM media containing low glucose concentration (LG; 8 mM). We used 8 mM glucose as the control condition since this concentration is representative of postprandial blood glucose levels in healthy individuals (≤7.8 mM) [15]. In addition, cells were also cultured in DMEM containing high glucose concentration (HG; 25 mM), with or without 50 µM boldine. When cultures reached ∼70% of confluence the media was replaced by differentiation media (DMEM 8 or 25 mM plus 5% horse serum) to promote the fusion of myoblasts and myotubes formation. Differentiation medium was renewed every 48 h, and boldine (50 µM) was added at each medium change.

### Evaluation of Etd^+^ uptake

It was carried out as previously described [16]. Briefly, myoblast plated onto plastic culture dishes were washed twice with Krebs solution. For time-lapse measurements, myoblasts were incubated in Krebs solution containing 5 μM Etd^+^. The intensity of Etd^+^ fluorescence was first recorded for 5 min in regions of interest by using a water immersion Olympus 51W1I upright microscope (Japan). Images were captured with a Retiga 13001 fast cooled monochromatic digital camera (12-bit; QImaging, Canada) every 15 s during 5 min, and image processing was performed offline with ImageJ software (National Institutes of Health).

### Intracellular Ca^2+^ signal

Intracellular Ca^2+^ signals were evaluated in immortalized human myoblast loaded with FURA 2. To load FURA 2, myoblasts were incubated with 5 µM FURA 2-AM in DMEM without serum at 37°C for 30 min and then washed three times. Then, the Ca^2+^ signals were evaluated using a Nikon Eclipse Ti microscope and fluorescence emitted upon excitation at two wavelengths (340 and 380 nm) was evaluated and the ratio of the recorded emissions (340/380 ratio), called Ca^2+^ signal, was calculated.

### Immunofluorescence analysis

Samples were fixed with 4% paraformaldehyde, incubated overnight at 4°C with diluted primary anti-PPAR-γ antibodies, followed by washes with PBS. Secondary antibodies conjugated to Cy2 or Cy3 were then applied for 1 hour. Samples were rinsed, mounted with fluoromount G (containing DAPI), and visualized on glass slides.

### Oil red O stain

Transverse cryosections of tibialis anterior muscle and cultured myoblasts were fixed with 4% paraformaldehyde in the presence of 180 mM CaCl₂. Oil Red O stock solution was diluted (3:2 with distilled water) to prepare the working solution, which was applied to samples for 30 min. Excess stains were removed, and samples were rinsed with water to visualize lipid droplets.

### Reverse transcription polymerase chain reaction (PCR)

Total RNA was isolated from tissue or cells using TRIzol following the manufacturer’s instructions (Invitrogen, Waltham, MA, USA). Two microgram aliquots of total RNA were transcribed to cDNA using MMLV-reverse transcriptase (Fermentas, USA), and mRNA levels were evaluated by PCR amplification (GoTaq Flexi DNA polymerase; Promega, USA)

The oligos used were the following: NLRP3: S 5’-GCTGGCATCTGGGGAAACCT-’3, AS 5’GCCCTTCTGGGGAGGATAGT-’3; CASP1 S 5’GAAAAGCCATGGCCGACAAG-’3, AS 5’-GCCCCTTTCGGAATAACGGA-’3. 18S: S 5′-TCAAGAACGAAAGTCGGAGG-′3, AS 5′-GGACATCTAAGGGCATCACA-′3.

### Statistical Analysis

Quantitative variables are presented as mean ± SEM. Considering data distribution, we used parametric or non-parametric tests, as appropriate, using the Shapiro–Wilk normality test. For multiple group comparisons during perfusion analysis, data are presented as mean values calculated from three different subjects per group. Multiple comparisons were performed using ordinary one-way ANOVA, followed by multiple comparisons by controlling the false discovery rate by using the Benjamini and Hochberg method. In other experiments, data were analyzed using one-way ANOVA followed by Tukey’s multiple-comparison test and an appropriate normality test. *p* < 0.05 was considered a statistically significant difference. Data and statistical analyses were performed using the Microsoft Excel database and GraphPad Prism 6 (GraphPad Software, La Jolla, CA, USA).

## RESULTS

### Boldine improves muscle strength and resting membrane potential in diabetic mice

To evaluate whether boldine preserves skeletal muscle function in diabetic mice, forelimb grip strength was measured at the end of the treatment period. STZ-treated mice exhibited significantly reduced grip strength compared to Control (Fig. 1A), consistent with diabetes-induced muscle weakness. Notably, boldine treatment prevented this functional decline and the values of grip strength were comparable to those observed in Control mice (Fig. 1A).

**Figure 1.**
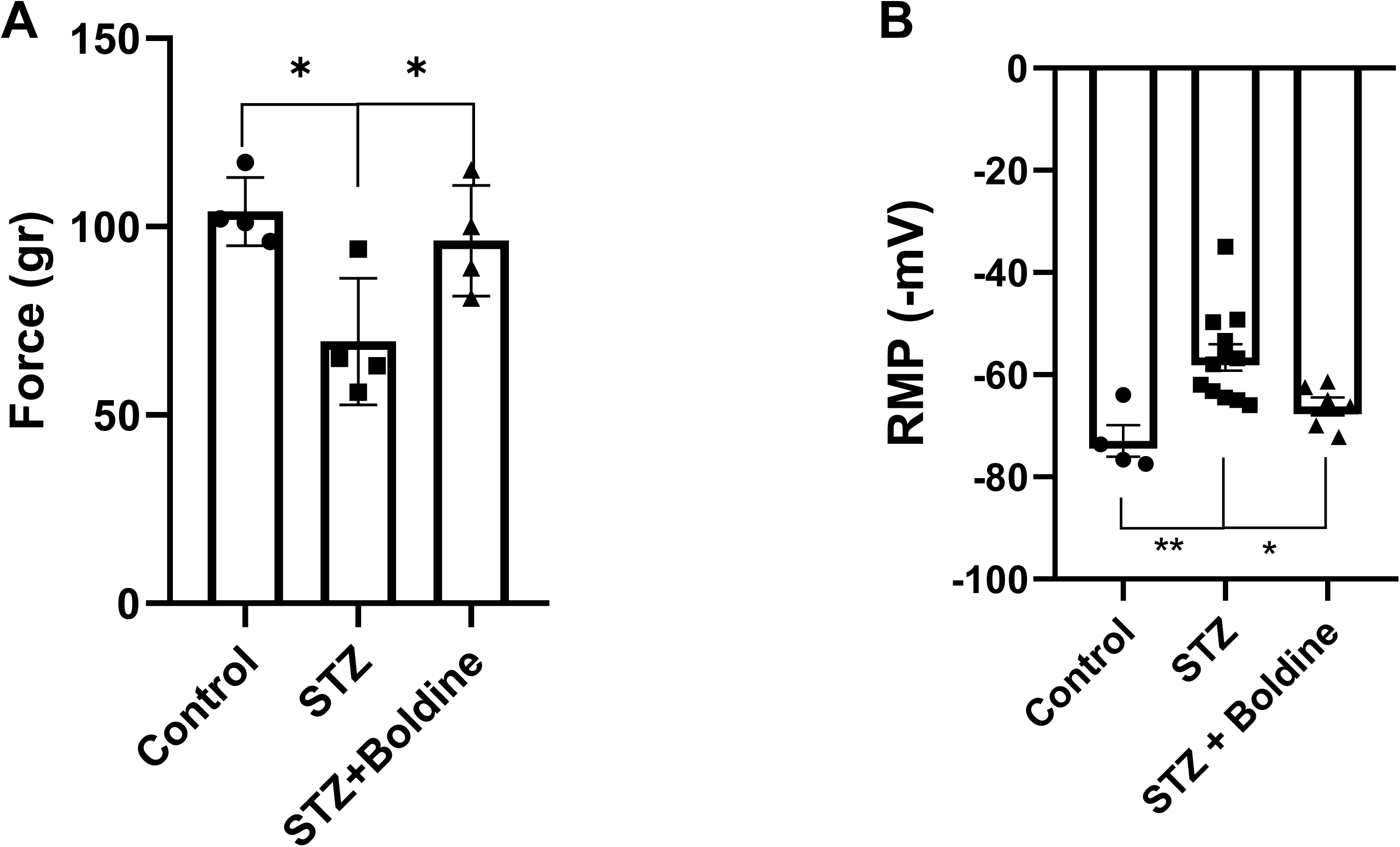
Boldine preserves forelimb grip strength and membrane potential in diabetic mice. **A)** Forelimb grip strength was measured using a force transducer before (Pre) and after (Post) the treatment period in wild-type (Control), STZ-induced diabetic (STZ), and diabetic mice treated with boldine (STZ + Boldine). STZ mice showed a reduction in grip strength following diabetes induction, whereas the force values of boldine-treated mice were comparable to their baseline measurements and to Control. **B)** Resting membrane potential (RMP) of isolated skeletal muscle fibers from Control, STZ, and STZ + Boldine mice. Fibers from STZ mice displayed significant depolarization compared to Control, which was prevented by boldine treatment. Data are presented as mean ± SEM. *p < 0.05, **p < 0.01, Tukey’s test.

To further assess muscle function at the cellular level, we measured RMP of freshly isolated skeletal muscle fibers. Fibers from STZ mice exhibited significant depolarization compared to Control fibers, which was prevented by boldine (Fig. 1B).

### Boldine maintains the acetylcholine-induced increase in blood perfusion

We next examined whether the loss of muscle strength in diabetic mice was associated with impaired endothelium-dependent microvascular perfusion, and whether boldine could restore this function. To do so, we assessed blood perfusion in the right gastrocnemius muscle (Figure 2A–2C).

**Figure 2.**
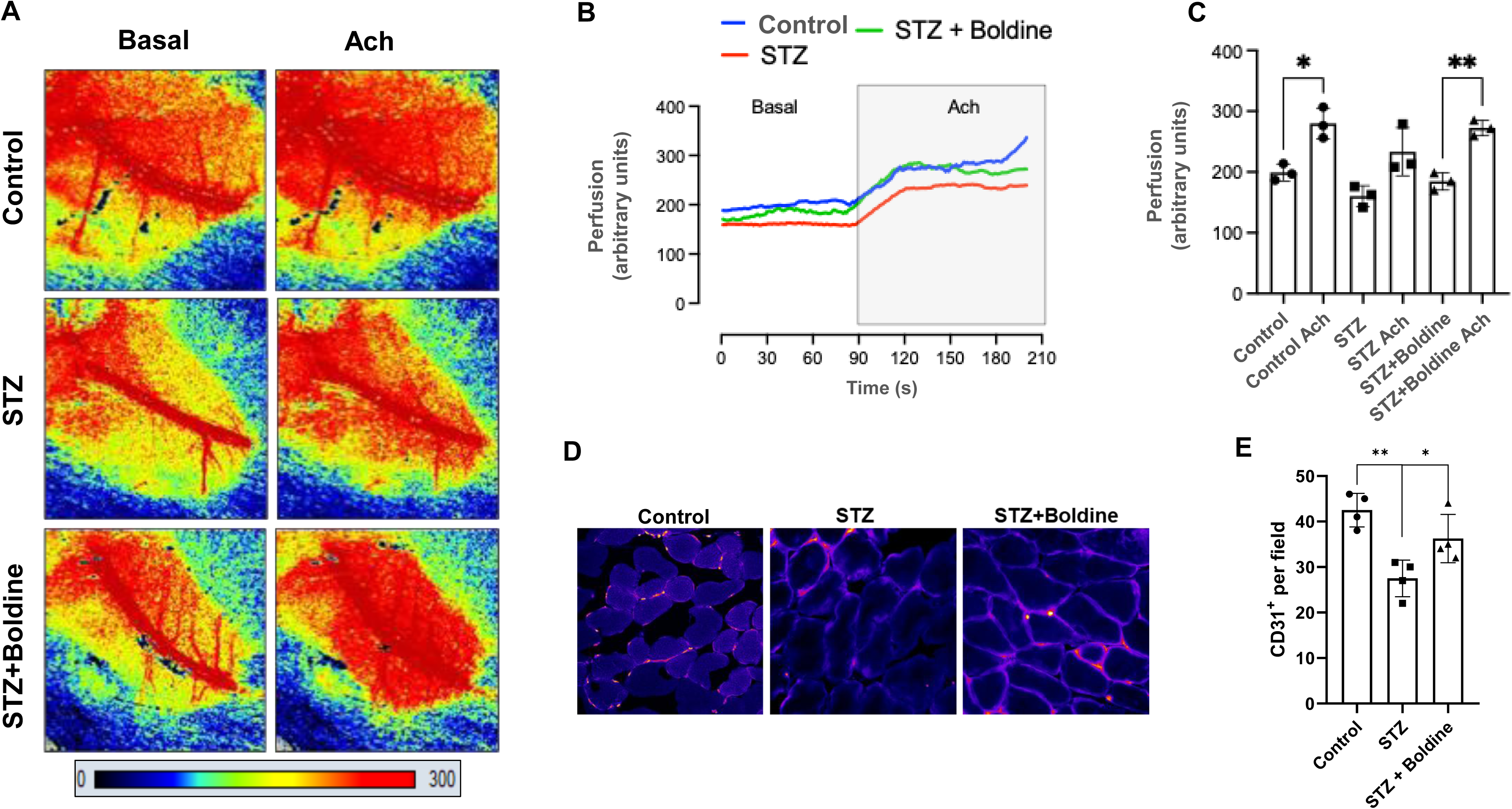
Blood perfusion of hindlimbs and microvasculature are preserved by boldine in diabetic mice. **A)** Representative images of blood perfusion in the right gastrocnemius muscle of wild-type (Control), STZ-induced diabetic mice (STZ), and boldine-treated STZ mice (STZ + Boldine) in absence (Basal) or presence of endothelial-mediated vasodilator acetylcholine (Ach, 10 μM). Perfusion units from 0 to 300, as indicated in the scale. **B)** Representative traces of blood perfusion as indicated in A. Highlighted area indicate perfusion after Ach stimulation. **C)** Average of perfusion unit in basal (Control, STZ and STZ + Boldine) conditions and same groups in presence of Ach. **D)** Representative immunofluorescence images of gastrocnemius muscle cross-sections from wild-type (Control), STZ-induced diabetic mice (STZ), and boldine-treated STZ mice (STZ + Boldine) stained for CD31 (red), an endothelial cell marker. Scale bar: 50 μm. **E)** Quantification of CD31⁺ capillaries per field shows a reduced capillary density in STZ-induced diabetic mice compared to Control, with recovery in boldine-treated STZ mice. Data are presented as mean ± SEM. n = 4.* p < 0.05, ** p < 0.01, Tukey test.

At baseline, myofibers of STZ-induced diabetic mice exhibited ∼20% lower microvascular perfusion expressed as units of perfusion compared to Control myofibers (160.1 ± 17.2 versus 199.1 ± 13.8 units, respectively, p=0.053). Notably, this decline was not observed in the STZ-induced diabetic mice treated with boldine (184.6 ± 14.3 perfusion units, p = 0.46).

We then evaluated the response to acetylcholine (ACh), an endothelium-dependent vasodilator. As expected, ACh elicited a robust increase in perfusion in Control mice that tend to be biphasic. In contrast, diabetic mice showed a blunted ACh-mediated response, denoting endothelial dysfunction. Boldine restored the increases in ACh-induced increase in perfusion (Figure 2C), which is in line with the improvement in muscle function (Figure 1A).

To investigate whether the boldine-induced improvement in endothelium-dependent tissue perfusion was accompanied by structural remodeling of the microvasculature, capillary density was assessed in gastrocnemius muscle sections by immunodetection of CD31, a specific marker of endothelial cells. Capillary-to-fiber ratio was used as an index of microvascular density and angiogenic capacity. STZ-induced diabetic mice exhibited a marked reduction in capillaries per muscle fiber compared with Control mice (Figure 2D, E), indicating an impaired skeletal muscle blood irrigation in the diabetic state. In contrast, boldine treatment significantly restored capillary density in diabetic muscles, as evidenced by an increased number of CD31⁺ structures per fiber relative to untreated STZ mice (Figure 2D, E). Collectively, these results indicate that boldine promotes microvascular remodeling in diabetic skeletal muscle and suggest that its beneficial effects on blood perfusion are associated with improved endothelial function and enhanced angiogenic responses.

### Boldine prevents lipid accumulation and up-regulation of NLRP3 mRNA levels in skeletal myofibers of diabetic mice

To determine whether boldine prevents the lipid accumulation observed in vitro in myoblasts exposed to high glucose, we analyzed cryosections of tibialis anterior muscle. Muscle sections were stained with Oil Red O to visualize neutral lipid droplets. In Control, virtually no Oil Red O–positive fibers were detected (Fig. 3, top panel). In contrast, myofibers of skeletal muscle from STZ-induced diabetic mice exhibited a marked increase in intracellular lipid accumulation (52.4 ± 3.6%), as evidenced by the presence of numerous red-stained myofibers (Fig. 3A). Boldine treatment markedly reduced intracellular lipid accumulation in diabetic mice (Fig. 3A) to a similar level to that observed in Control (Boldine: 15.2 ± 4.1% and Control: 3.1 ± 1.3%, p > 0.05). These results indicate that boldine prevents lipid infiltration into skeletal muscle fibers in diabetic mice.

**Figure 3.**
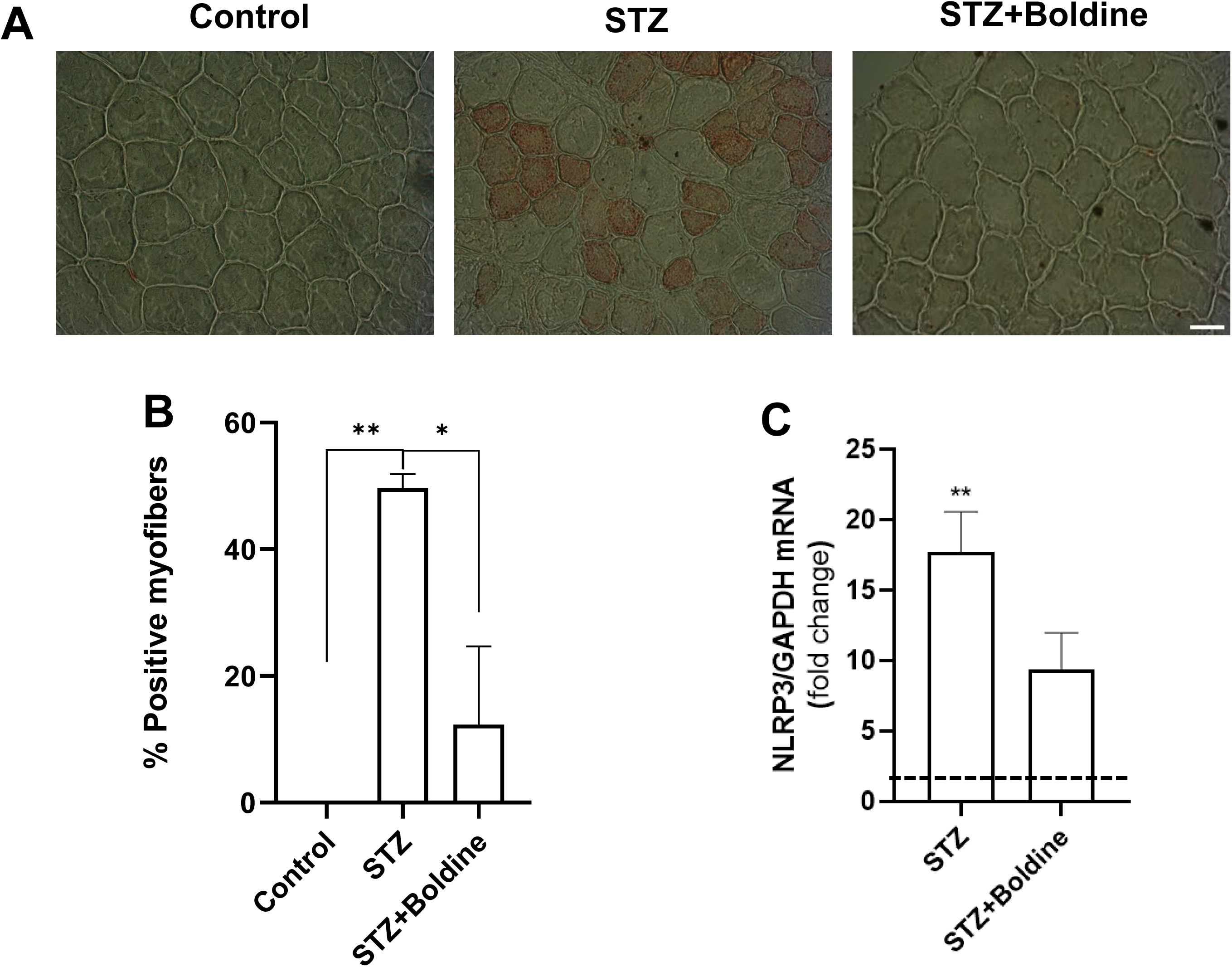
Boldine prevents lipid accumulation and increase in NLRP3 mRNA levels in skeletal muscle of diabetic mice. **A)** Representative images of tibialis anterior cryosections stained with Oil Red O from wild-type (Control), streptozotocin-induced diabetic (STZ), and diabetic mice treated with Boldine (STZ + Boldine). Lipid accumulation is evident in the STZ group, while minimal staining is observed in Control and STZ + Boldine mice. Scale bar: 50 µm. **B)** Quantification of the percentage of Oil Red O–positive myofibers per field. Data represent mean ± SEM (n = 4 mice per group). *p < 0.05, **p < 0.01, Tukey’s test. **C)** NLRP3 were evaluated by qPCR. n=4. **p<0.01 vs Control, Tukey test.

The inflammasome in skeletal myofibers has been shown to be activated by hyperglycemia [16]. To determine whether this alteration is also present *in vivo*, we quantified NLRP3 mRNA expression in skeletal muscle from diabetic mice by qPCR. Skeletal muscle from STZ-treated mice exhibited a marked upregulation of NLRP3 expression (17.74 ± 2.81-fold vs. Control; Fig. 3C), consistent with inflammasome activation under diabetic conditions. Boldine treatment significantly attenuated NLRP3 overexpression, reducing transcript levels by approximately 50%. However, NLRP3 expression remained significantly elevated compared with Controls (9.41 ± 2.56-fold vs. Control, p < 0.05). Together, these findings indicate that diabetes induces inflammasome activation in skeletal muscle, which is partially reversed by boldine.

### High glucose induces an increase in membrane permeability through large-pore channels

In a previous study, we found that HG increases the membrane permeability to Etd^+^ in myofibers [16]. Now, we have investigated whether HG could increase the activity of large-pore channels in myoblasts. Cells were cultured with medium containing LG (8 mM), HG (25 mM), or HG plus 50 µM boldine.

Cell membranes show low Etd⁺ permeability due to minimal hemichannel opening. Glucose at 8 mM within postprandial levels does not alter myoblast permeability or differentiation [15]. In contrast, pathological hemichannel upregulation increases Etd⁺ uptake, which becomes fluorescent upon nucleic acid binding.

In this study, we found that compared to LG (control condition), HG increases the fluorescence intensity over time, reflecting progressive Etd^+^ uptake (Fig. 4A and B). Notably, boldine prevents elevated Etd^+^ uptake generated by HG alone (Fig. 4A and B). Accordingly, cells differentiated under HG exhibited a significantly increased Etd⁺ uptake rate compared with those maintained in LG or HG supplemented with boldine (Fig. 4C). Importantly, this effect was not attributable to hyperosmolarity, as exposure to equiosmolar mannitol (25 mM) did not alter Etd⁺ uptake (Fig. 4C).

**Figure 4.**
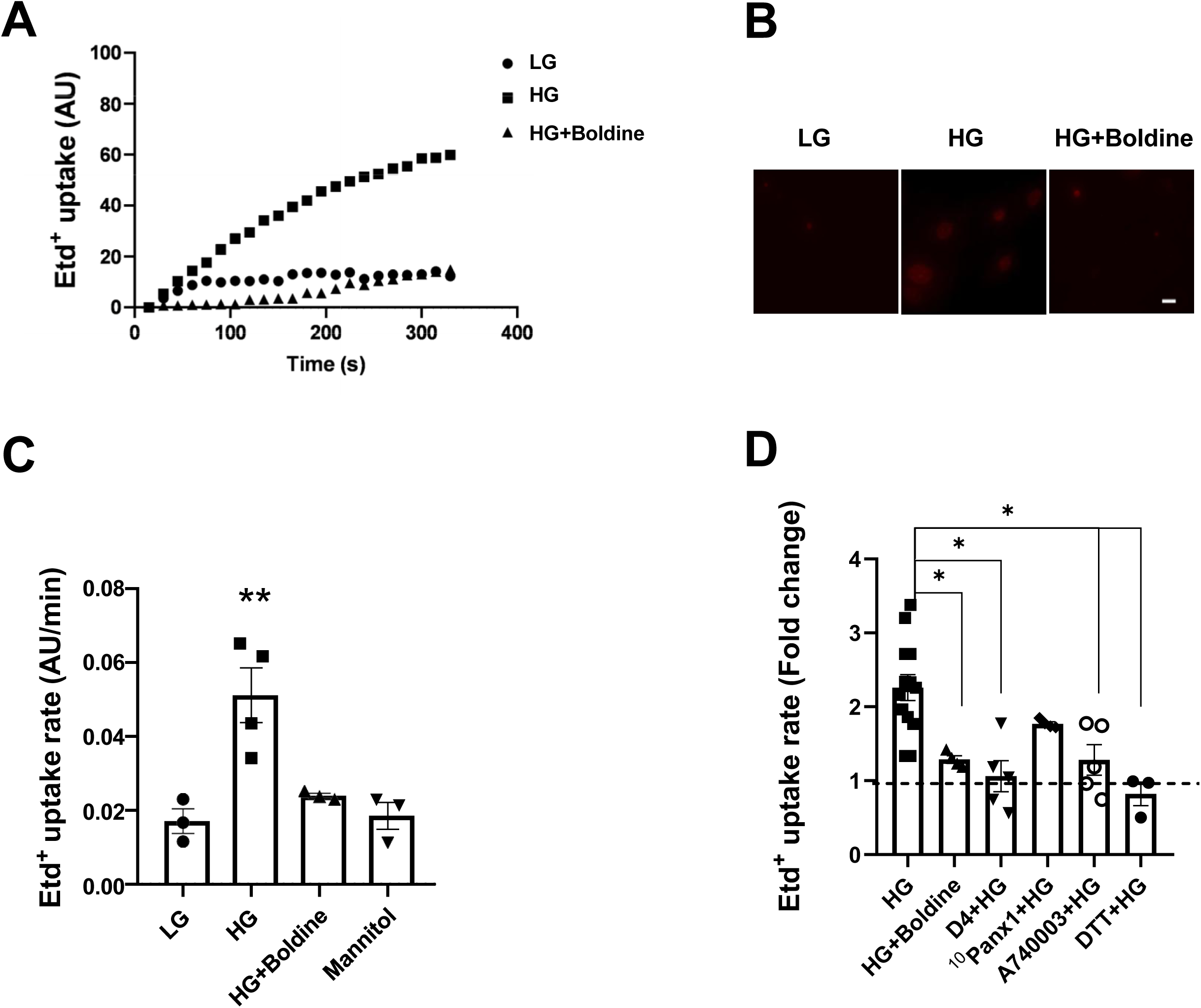
Differentiation to muscle fate in high glucose increases the membrane permeability of myoblasts. **A)** The Etd^+^ uptake assay was performed in myoblast treated with 8 mM glucose (LG), 25 mM glucose (HG), HG plus 50 µM boldine, or LG plus 25 mM Mannitol. The graph represents the fluorescence intensity of Etd^+^ uptake over time. Each point represents the average of ≥15 cells analyzed. **B)** Representative photographs of Etd^+^ fluorescence (Red) view of a field at the end of an experiment in each condition. **C)** Etd^+^ uptake rate of myoblasts treated with different conditions. **p<0.01, Tukeýs Test. Data were obtained from four independent experiments with four repeats each one (≥15 cells analyzed for each repeat). **D)** After 7 days, dye uptake was performed with different blockers: 50 µM boldine (Cx and Panx1 HCs); 100 nM D4 (Cx HC blocker); 200 µM ^10^Panx1 peptide (Panx1 HC blocker); 10 µM A740003 (P2X7R blocker)

To define the contribution of candidate large-pore channels to Etd⁺ uptake, we employed pharmacological inhibitors targeting HCs and the P2X7 receptor (P2X7R). We used D4, which selectively blocks of Cx45 and Cx43 HCs [14], and A740003, a selective P2X7R antagonist [17]. Both inhibitors completely abolished the HG-induced increase in Etd⁺ uptake in myoblasts (Fig. 4D), demonstrating the requirement of Cx HCs and P2X7R signaling in this response. In contrast, acute inhibition of Panx1 hemichannels with ^10^Panx1 [18] did not result in a significant reduction of Etd⁺ uptake (Fig. 4D), arguing against a major role for Panx1 channels. Collectively, these results indicate that the HG-induced increase in membrane permeability is mediated predominantly by connexin hemichannels rather than Panx1.

### High glucose activates connexin hemichannels through elevated intracellular Ca²⁺ and NO-dependent mechanisms

Activation of Cx43 HCs and P2X7Rs has been linked to intracellular Ca²⁺ elevation and purinergic signaling [19,20]. Given that increased intracellular Ca²⁺ enhances Cx43 HC open probability [21], we tested whether Ca²⁺ mobilization from intracellular stores contributes to the HG-induced increase in membrane permeability. Myoblasts cultured under HG conditions were preincubated for 5 min with MRS2179 (10 µM), a selective P2Y1 receptor antagonist that prevents IP₃-mediated Ca²⁺ release [22]. Although MRS2179 tended to reduce Etd⁺ uptake, this effect did not reach statistical significance (Fig. 5A), indicating that P2Y1-mediated Ca²⁺ mobilization from intracellular stores does not play a major role in HG-induced membrane permeabilization. Previous studies have demonstrated that p38 MAPK enhances Cx43 HC activity in response to pro-inflammatory stimuli [23]. To assess the acute involvement of this pathway in HC regulation while minimizing secondary signaling effects, cells were preincubated for only 5 min with pharmacological inhibitors prior to Etd⁺ uptake measurements. Inhibition of p38 MAPK with SB202190 (10 µM) reduced Etd⁺ uptake to levels comparable to those observed under LG conditions (Fig. 5A), implicating p38 MAPK signaling in HG-induced membrane permeabilization.

**Figure 5.**
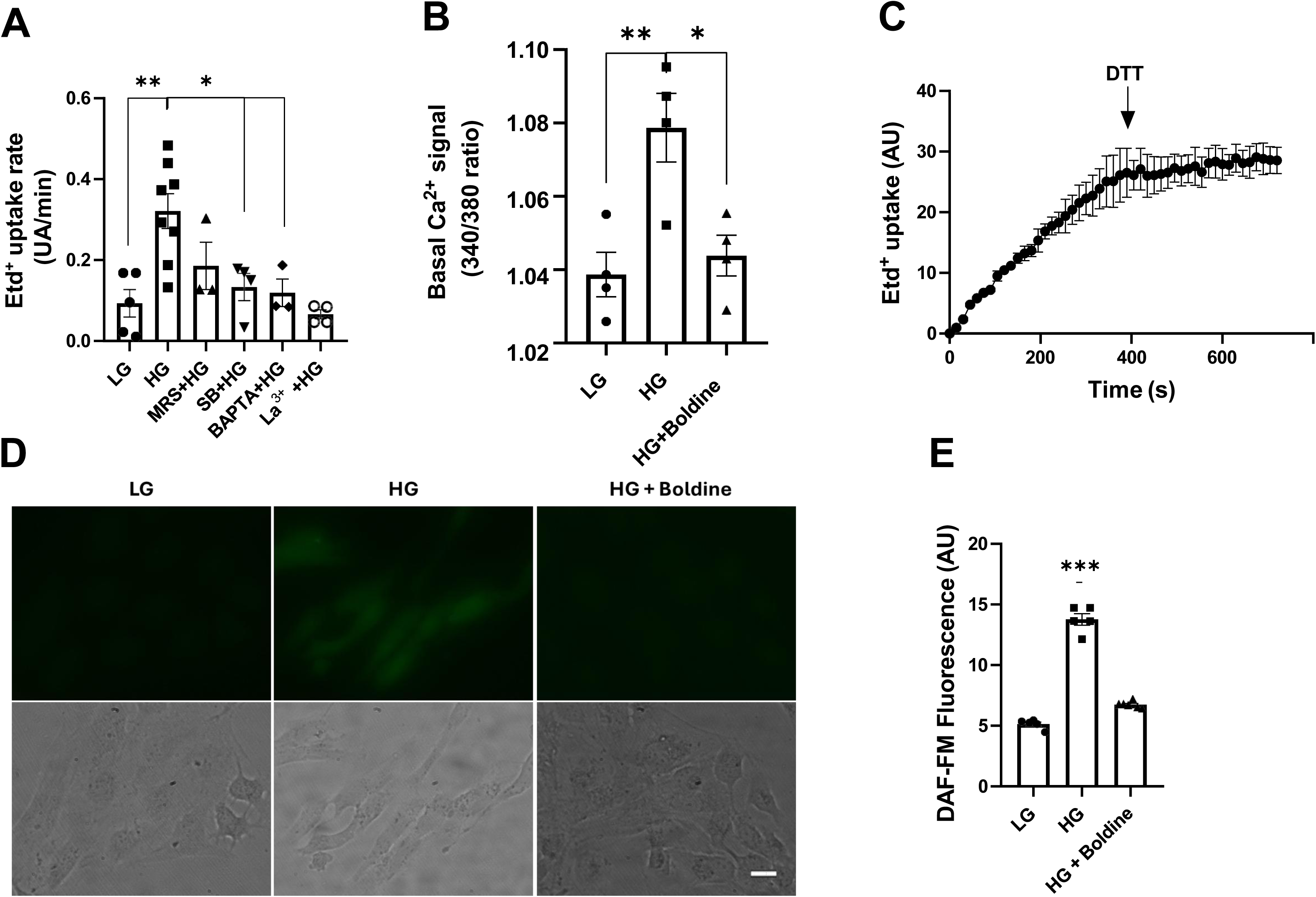
High glucose increases basal cytoplasmic Ca^2+^ signal and generates high levels of nitric oxide. Myoblasts (AB1167) were induced to acquire commitment for muscle differentiation in different concentrations of extracellular glucose. After 7 days, dye uptake was performed. **A)** The Etd^+^ assay was performed in myoblast treated with HG alone or in myoblasts preincubated for 5 min with the following agents: 10 µM SB202190 (p38), 10 µM MRS2179 (P2Y1R), or 10 mM DTT (Cystine reducer). Some myoblast cultures were loaded with 10 µM BAPTA-AM (BAPTA: Ca^2+^ chelator) before evaluating Etd^+^ uptake. The effect of 200 µM La^3+^ on the uptake of myoblasts culture in HG (La^3+^ + HG) was also evaluated. n=4. *p<0.05, **p<0.01 Tukeýs Test. Data were obtained from four independent experiments with four repeats each one (≥15 cells analyzed for each repeat). **B) .** Ca^2+^ signal denoted as Fura-2 ratio (340/380 nm excitation) of myoblast treated with different glucose concentrations. B) Basal intracellular Ca²⁺ signal (Fura-2 ratio, F/F₀) in cells treated with low glucose (LG), high glucose (HG) or HG plus 50 µM boldine (HG + Boldine). (n=4 with ≥15 cells recorder in each case). *p<0.05, **p<0.01 Tukeýs Test. **C)** Timelapse of Etd^+^ uptake in myoblast treated with HG, before and after application of 10 mM DTT (arrow). **D)** Representative micrographs of the basal level of nitric oxide (NO) in myoblast cultured in each condition. **E)** Bar graph representing the average of DAF fluorescence by myoblasts cells treated as mentioned above. n=3. ***p<0.001, Tukeýs Test. (≥15 cells analyzed for each repeat).

To further determine the contribution of intracellular Ca²⁺ signaling, myoblasts were loaded with BAPTA-AM (5 µM) to chelate cytosolic Ca²⁺. This intervention significantly reduced Etd⁺ uptake (Fig. 5A), demonstrating that elevated intracellular Ca²⁺ is required for HG-induced increases in the membrane permeability. Consistently, La³⁺, a broad inhibitor of Cx HCs and P2X7R [24], normalized Etd⁺ uptake in HG-treated cells up to LG levels (Fig. 5A), further supporting a role for large-pore channels in mediating hyperglycemia-induced permeabilization.

Intracellular Ca²⁺ dynamics were then directly assessed using Fura-2 AM in myoblasts differentiated under LG, HG, or HG supplemented with boldine (Fig. 5B). Cells exposed to HG displayed a significantly elevated basal Ca²⁺ signal relative to LG controls, whereas boldine fully prevented this increase (Fig. 5B). Quantitatively, the mean basal Ca²⁺ was highest in the HG group (1.0800 ± 0.0036), while values in LG and HG + Boldine were comparable (1.030 ± 0.005 and 1.040 ± 0.002, respectively) and significantly lower than HG alone.

Increased cytoplasmic Ca²⁺ levels are known to promote oxidative and nitrosative stress through the generation of reactive oxygen and nitrogen species [25]. In addition, S-nitrosylation of Cx43 [26] in cysteine residue 271 [27] has been shown to increase Cx HC activity. To evaluate whether redox-dependent regulation contributes to HC activation during myoblast differentiation under HG-conditions, cells were treated with the reducing agent DTT (10 mM). Acute application of DTT rapidly and significantly decreased Etd⁺ uptake, reaching a stable minimum within seconds (Fig. 5C), consistent with reduced membrane permeability. These results indicate that Cx43 HC opening under HG involves redox-sensitive mechanisms.

Given that HG increases nitric oxide synthase expression and nitric oxide (NO) production in cell cultures [28], we tested whether AB1167 cells treated with HG presented an increased NO production. We found a significant increase in NO levels in cells cultured in HG, but not in cells incubated in LG or HG plus boldine (Fig. 5D and E). These results suggest that S-nitrosylation of Cx43 HCs might contribute to increased Edt^+^ uptake induced by HG.

### Boldine prevents the increase in mRNA levels of inflammasome components triggered by high glucose

Because HG is known to activate the inflammasome in skeletal muscle cells [16], we examined whether myoblasts differentiated under HG conditions for 7 days showed evidence of inflammasome activation. Cells differentiated in HG exhibited increased mRNA levels of NLRP3 and caspase-1 compared with cells differentiated in LG (Fig. 6A, B). These findings indicate that HG promotes inflammasome-related gene expression during myoblast differentiation. We further propose that the increased membrane permeability to Etd⁺ observed under HG conditions may contribute to this inflammatory response.

**Figure 6.**
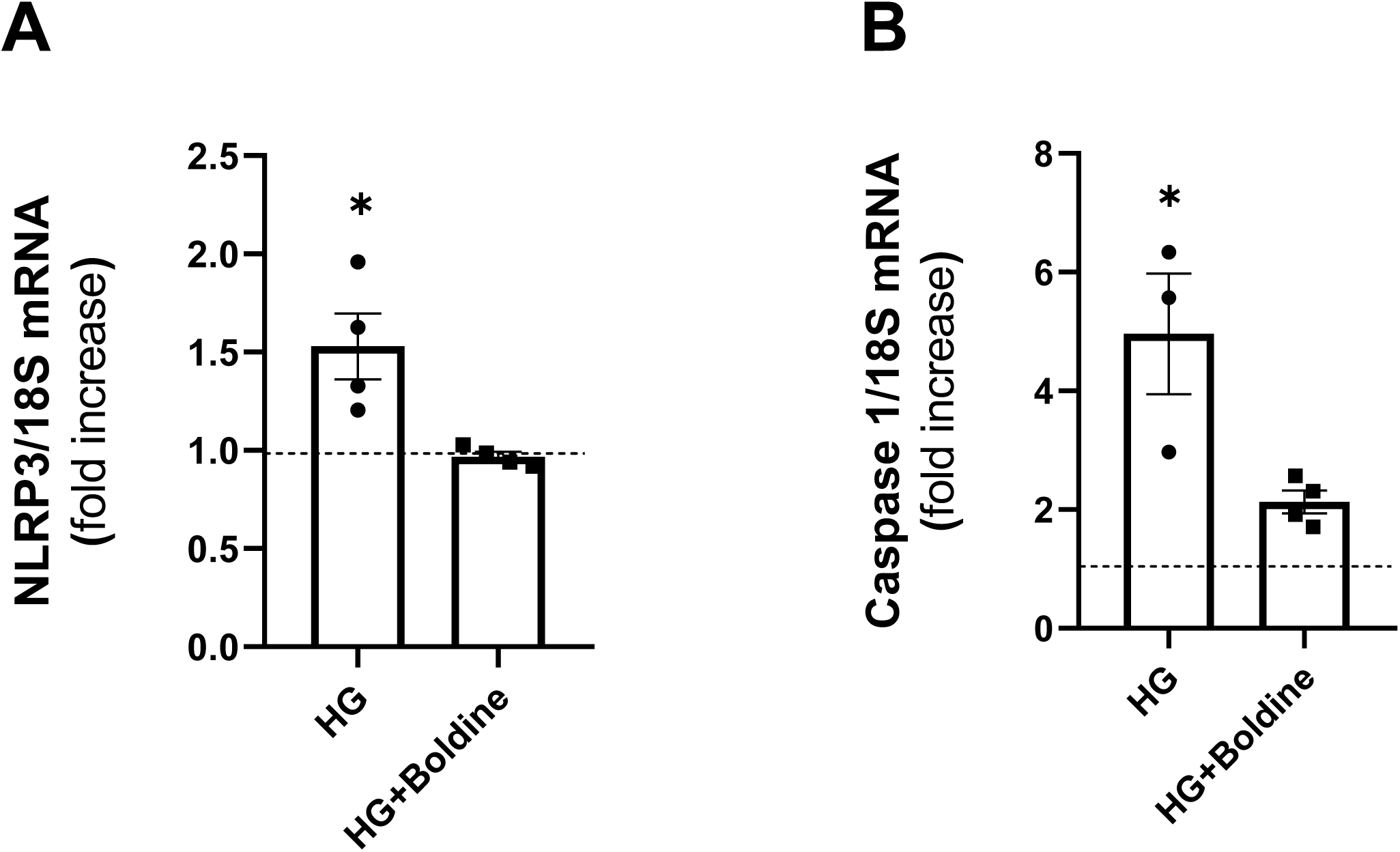
Boldine prevents the increase in mRNA levels of inflammasome components. Myoblasts were cultured in low (LG) high (HG) glucose concentration (8 and 25 mM, respectively), or HG plus boldine (HG + Boldine) in medium that induces acquisition of skeletal muscle commitment. After 7 days in culture the mRNA levels of **A)** NLRP3, and **B)** caspase-1 were evaluated by qPCR. n=3. *p<0.05 vs LG, Tukey test. Dashed lines indicate the mean of control values.

### Boldine prevents high-glucose–induced adipogenic drift during myogenic differentiation

Skeletal muscles of adult diabetic men show intramuscular triglycerides accumulation [29], which could result in reduction of muscular strength, as observed in other pathologies [12]. Here, we evaluated if immortalized human myoblasts exposed to HG could differentiate to adipocytes as a possible explanation of intramuscular fat accumulation [29].

After 7 days of differentiation, we found that HG elevated the nuclear content of the adipogenic transcription factor PPARγ; where ∼45% presented nuclear PPARγ reactivity (Fig. 7A and B). Notably, this response was prevented by boldine treatment (50 µM) (Fig. 7A and B). In addition, we observed the presence of triglycerides accumulation in several myoblasts cultured in differentiation media with HG (red droplets). Boldine treatment prevented this phenomenon (Fig. 7C). Thus, boldine blocked the aberrant differentiation of myoblasts to fat-containing cells.

**Figure 7.**
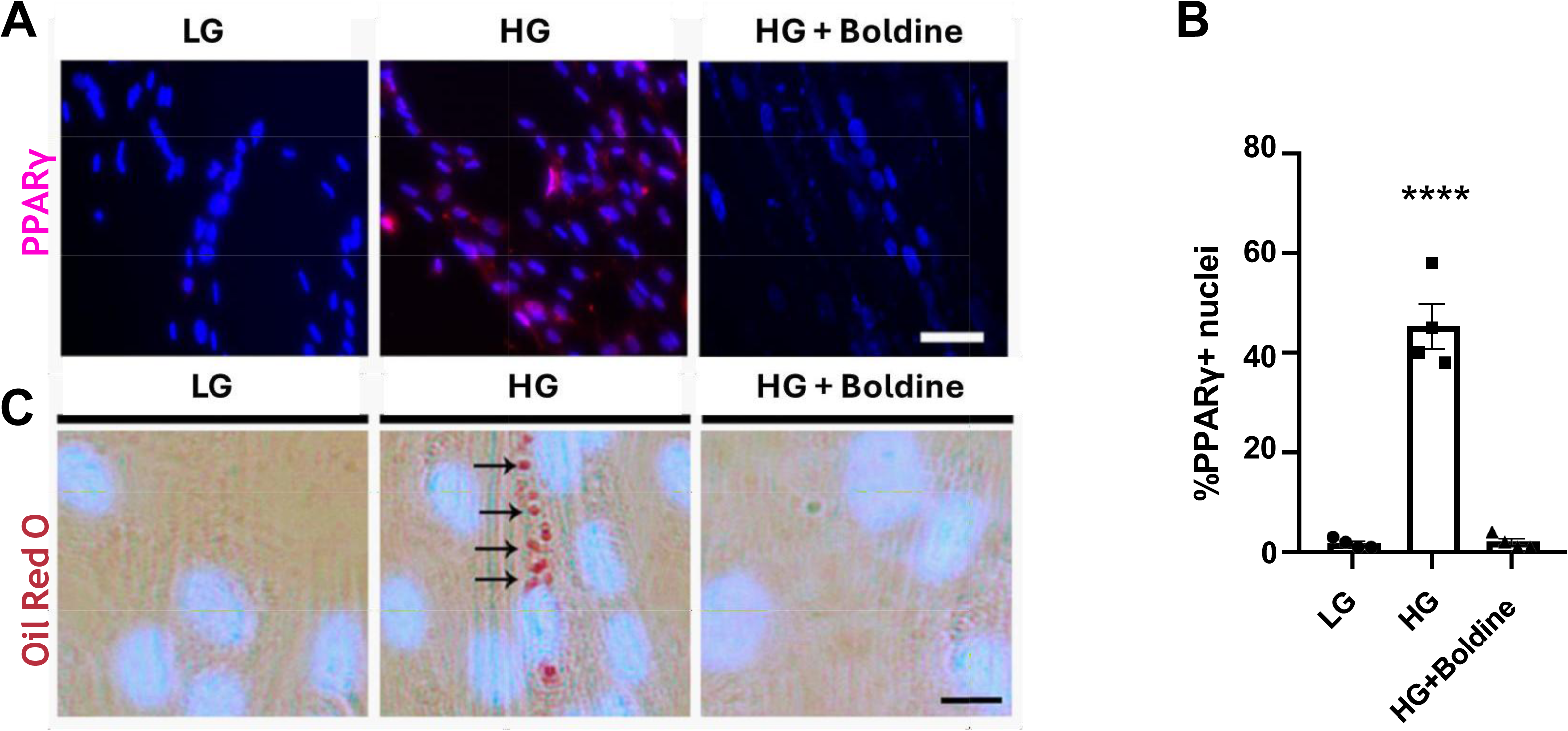
Boldine prevents the acquisition of adipogenic commitment and differentiation to adipocyte-like cells of myoblast induced to acquire myogenic commitment in high glucose. Myoblasts were incubated in low (LG) and high glucose (HG) conditions (8 and 25 mM, respectively) in the medium that induces differentiation with myogenic differentiation medium in HG plus 50 µM boldine (HG + Boldine). After 7 days, they were fixed for immunofluorescence. **A)** Analysis against PPAR (fuchsia) and nuclear staining with DAPI (blue). n=4 cell cultures. **B)** Quantification of PPARγ positive nuclei expressed as percentage (% PPARγ + nuclei) from 10 fields like in (A). n=4 cell cultures. ****p<0.0001, Tukeýs Test. **C)** Detection of triglyceride accumulation with oil Red O staining (arrow). n=4 cell cultures. Scale bar: 10 µm. **D)** Microphotograph showing cells positive for oil Red O staining. n=4 cell cultures. ****p<0.0001, Tukeýs Test.

## DISCUSSION

STZ treatment increased HC activity in skeletal muscle, accompanied by higher levels of inflammasome components, lipid accumulation, reduced muscle force, and impaired blood perfusion. High glucose similarly increased HC activity in myoblasts, elevated the mRNA levels of inflammasome components, and promoted lipid accumulation. Boldine prevented these alterations in both models and preserved muscle function.

Diabetes-induced skeletal muscle weakness is consistently observed in rodent models [30], an effect that was confirmed in this study showing reduced forelimb grip strength in STZ-induced diabetic mice.

Importantly, boldine treatment fully prevented this decline. This functional impairment is closely associated with a significant depolarization of RMP in skeletal muscle fibers from diabetic mice, an indicative of altered membrane excitability. Such RMP depolarization has also been described for other muscle pathologies, such as denervation [14] and sepsis [31], where connexin HCs contribute to excitability loss. Early studies similarly noted reduced RMP in genetically diabetic and alloxan-induced diabetic mice, with these changes correlating with blood glucose levels and disease duration [32]. Our group previously demonstrated that diabetes induces *de novo* expression and membrane redistribution of Cx39, Cx43, and Cx45 in skeletal muscle, a pathological pattern that boldine treatment effectively reversed [16]. Crucially, genetic deletion of Cx43 and Cx45 in skeletal muscle prevents significantly diabetes-induced muscle atrophy [16], strongly implicating Cx HCs in both the structural and functional decline of skeletal muscle under hyperglycemic conditions. Our current findings reinforce this evidence, showing that boldine not only restores RMP and contractile strength but also preserves muscle mass and excitability by inhibiting HC activity, thereby targeting a unifying mechanism underlying diabetic myopathy.

Beyond reduced contractility and excitability, diabetic muscle displays a marked inflammatory profile, particularly involving inflammasome activation. HG glucose-mediated inflammasome activation is well-documented in diabetic patients, who exhibit elevated plasma levels of IL-1β and IL-18. In vitro studies, including our previous report, show increased NLRP3 levels in skeletal myofibers treated with HG [16]. *In vivo* studies in STZ-induced diabetic mice and db/db mice have also demonstrated upregulation of NLRP3 and caspase-1 [33]. Our data further demonstrate that blocking connexin HC with boldine prevents the increase in mRNA levels of NLRP3 and caspase-1, suggesting a direct involvement of connexin HC in inflammasome activation in this context.

Another prominent hallmark of diabetic myopathy is intramuscular lipid accumulation. Consistent with previous findings in diabetic rats [29], the STZ-induced diabetic mice used in the present work exhibited a marked lipid accumulation in tibialis anterior muscle fibers. Notably, boldine treatment significantly reduced the number of lipid-positive myofibers in diabetic mice, reaching levels observed in non-diabetic Controls. This effect aligns with prior work in a dysferlinopathy model, where boldine administration prevented intramuscular fat accumulation and reduced PPARγ expression [12]. Our *in vitro* findings further demonstrate that boldine prevents HG–induced lipid accumulation in human myoblasts undergoing myogenic differentiation, extending its protective role against pathological ectopic fat deposition. In parallel, the restoration of muscle perfusion and microvascular network organization observed *in vivo* in boldine-treated diabetic mice highlights its systemic vasoprotective actions, which likely contributes to improve muscle integrity and functional capacity.

To understand the molecular mechanisms, our study revealed that HG induces an aberrant adipogenic commitment in myoblasts, a process found to involve large-pore channel activation by a mechanism sensitive to boldine [34]. Boldine prevented the HG-induced increase in myoblast membrane permeability, an effect also observed with acute boldine pre-incubation, indicating a direct channel-blocking action. By using more specific blockers, we identified Cx HCs and P2X7Rs as the main mediators of this permeability increase. In contrast, blocking Panx1 HCs and P2Y1Rs caused non-significant effects. Likely, P2Y1Rs undergo desensitization under conditions of sustained extracellular ATP release [35], which may explain these results. These results support a positive feedback mechanism in which ATP released through Cx HCs activates P2X7Rs, promoting Ca²⁺ entry. Since both P2X7Rs and Cx43 HCs are permeable to Ca²⁺ [20,36], this rise in intracellular Ca²⁺ further enhances Cx43 HC opening [21].

In line with the critical role of intracellular free-Ca²⁺, HG significantly increased basal cytoplasmic Ca²⁺ signals, which were effectively reduced by the Ca²⁺ chelator BAPTA. Additionally, p38 MAPK activation, which previously was linked to Cx43 HC activity in inflammatory contexts, was found to increase Etd^+^ uptake in human myoblasts [23]. Furthermore, high NO production in diabetes, caused by HG-induced upregulation of NO synthase [28], explains the elevated NO levels detected in HG-exposed myoblasts. NO enhances Cx43 HC activity through S-nitrosylation of cysteine residues [26,27], and the reversal of HG-induced permeability by DTT supports the involvement of S-nitrosylated Cx43 HCs.

Elevated Cx HC activity also contributes to increased reactive oxygen species (ROS) production, which can be mitigated by HC inhibitors or antioxidants. Since Cx43 HCs are permeable to Ca²⁺ [37], blocking these channels may prevent activation of Ca²⁺-dependent pathways responsible for ROS generation. Boldine possesses intrinsic antioxidant properties [34], suggesting that its dual action (i.e., blockade of Cx HCs and reduction of oxidative stress) offers a comprehensive therapeutic approach to diabetes-related myopathy.

HG-induced adipogenic differentiation of muscle-derived stem cells may be directly linked to this increased ROS production [5]. Elevated oxidative stress has been shown to transform myoblasts into brown adipocytes by reducing MyoD expression via NF-κB activation [38]. Our findings that boldine prevented HG-induced increases in PPARγ levels and lipid accumulation in myoblasts are consistent with this mechanism. Given that boldine significantly reduces fat infiltration in dysferlinopathy models [39], we propose that connexin HC represents a common therapeutic target to limit fat infiltration in muscle diseases characterized by metabolic or inflammatory stress, including diabetes, obesity, dysferlinopathy and Duchenne muscular dystrophy [6].

In conclusion, our data identify large-pore channels as key drivers of diabetes-associated skeletal muscle dysfunction and demonstrate that boldine restores muscle function, vascular reactivity, and metabolic and inflammatory homeostasis *in vivo* and *in vitro*. These findings highlight the potential efficacy of large-pore channel inhibition as a disease-modifying strategy and position boldine as a promising therapeutic candidate for diabetic myopathy.

## Acknowledgments

This work was partially funded by ANID grant 1231523 and ICN2025_026 CINV (to J.C.S.) and Doctoral fellowship grants from ANID 21211179 (W.V.) and 21241885 (A.L.). The experimental data in this paper were drawn from a thesis submitted in partial fulfillment of the requirements for the Doctorate in Biological Sciences, with a mention in Physiological Science (W.V.) at Pontificia Universidad Católica de Chile. C.E. is funded by Fondecyt grant 1240295 and GI2301146.

## Conflict of interest

The authors declare no conflict of interest.

Use of AI Tools

Portions of this manuscript were written with OpenAI’s ChatGPT (GPT-4, July 2025 version) to improve clarity, language, and coherence. The authors critically reviewed, edited, and approved all AI-assisted content. No AI tools were used for data analysis, result interpretation, or the generation of scientific conclusions.

## Data Availability

All data supporting the findings of this study are included in the manuscript and its Supporting Information files. Additional datasets are available from the corresponding author, Prof. Juan C. Sáez, upon reasonable request and subject to institutional and ethical approval.

